# Intermittent Cocaine Use Patterns, Not Total Intake, Predict Cue-induced Drug Seeking in Rats

**DOI:** 10.1101/2025.10.21.683821

**Authors:** Amélie Mainville-Berthiaume, Hajer Algallal, Salomé Dupuis, Sema Abu Shamleh, Coralie Godbout, Vincent Jacquemet, Anne-Noël Samaha

## Abstract

**Background and purpose:** Cocaine-associated cues trigger relapse to drug use in humans and animal models. In rats, long daily cocaine access (4-6 h vs. 1-2 h) increases drug self-administration and cue-induced cocaine seeking, suggesting that greater intake promotes relapse. However, prior studies used continuous-access paradigms, whereas human cocaine use is typically intermittent. Moreover, these studies used conditioned stimuli (CS), whereas discriminative stimuli (DS) are more effective in triggering increases in drug-seeking actions. Here, we used intermittent access (IntA) self-administration to examine how session length (Short-IntA: 2 h vs. Long-IntA: 4 h) affects CS- and DS-induced cocaine seeking.

**Experimental approach:** Female rats self-administered cocaine intermittently during daily 2- or 4-h IntA sessions, with alternating DS+ (cocaine available; 5 min) and DS- (no cocaine; 25 min) periods. Lever pressing during DS+ delivered cocaine and a CS+; lever pressing during DS-delivered only a CS-. After four weeks of abstinence, rats received a cocaine-seeking test where all cues were presented response-independently, and lever presses - which had no consequence - measured cocaine seeking.

**Key results:** Long-IntA rats took twice more cocaine than did Short-IntA rats. Both groups later showed significant (DS+)-but not (CS+)-triggered cocaine seeking, with no group differences. Thus, total intake did not predict cue-induced relapse propensity. However, individual cocaine intake patterns did (hourly consumption, burst-like self-administration, and latency to first infusion).

**Conclusions and Implications:** Under intermittent-access conditions, individual cocaine-use patterns, not cumulative intake, predict vulnerability to cue-induced relapse. This highlights the importance of individual drug-taking profiles in relapse risk assessment.

## Introduction

Preventing relapse to cocaine use following periods of abstinence is a major challenge in treating individuals with cocaine use disorders (O’Brien & Gardner, 2005). This is in part because environmental stimuli linked to past cocaine use (e.g., people, places and objects) are very effective in triggering drug cravings and relapse in cocaine users (O’Brien et al., 1992; Vafaie & Kober, 2022). Similarly, cocaine-associated cues evoke, sustain and invigorate cocaine-seeking actions in laboratory rats (Stewart et al., 1984; for review see Shaham et al., 2003).

More extensive cocaine self-administration is thought to increase vulnerability to cue-induced relapse. Rats with Long-Access to cocaine (6 h/session) show greater cue-induced cocaine-seeking behaviours compared to those with Short-Access (1-2 h/day) (Ferrario et al., 2005; Fischer et al., 2013; Jin et al., 2010; Kippin et al., 2006). Long-Access promotes high and escalating levels of cocaine self-administration compared to Short-Access, where intake levels often remain low and stable across sessions (Ahmed et al., 2000; Ahmed & Koob, 1998). Consequently, a common assumption is that “*excessive drug exposure likely remains an indispensable element driving the development of addiction*” and that “*addiction-causing neuropathological processes could be set in motion only when rats can expose themselves sufficiently to cocaine to cross the ‘threshold of addiction’*” (Ahmed, 2012).

Short- and Long-Access procedures involve continuous access to cocaine during each self-administration session, and this produces sustained brain cocaine concentrations throughout each session (Algallal et al., 2020; Allain et al., 2018; Zimmer et al., 2012). However, human cocaine use is typically intermittent, both within and between bouts of intake (Beveridge et al., 2012; Cohen & Sas, 1994; Leri et al., 2004), producing a spiking pattern of brain cocaine concentrations. Intermittent access (IntA) self-administration models this pattern in rats by providing periods of cocaine availability (e.g., 5 min) interspersed with periods of cocaine unavailability (e.g., 25 min) during each self-administration session. Although IntA procedures result in less total cocaine intake per session compared to Long-Access, with intake levels similar to continuous Short-Access, IntA produces stronger addiction-like features (Allain et al., 2015; Kawa et al., 2019; Samaha et al., 2021). These include greater and more persistent motivation for cocaine (Allain et al., 2018; James et al., 2019), continued drug taking despite adverse consequences (James et al., 2019) or drug unavailability (Kawa et al., 2016), and more robust cue-induced cocaine seeking behaviour (James et al., 2019; Kawa et al., 2016; Nicolas et al., 2019).

Findings from IntA studies have challenged the idea that taking large, escalating amounts of cocaine is necessary to produce addiction-like features in rats. Shorter IntA cocaine self-administration sessions (2 h/session) are just as effective as longer IntA sessions (6 h) in promoting psychomotor sensitization, incentive motivation for cocaine and cocaine-primed reinstatement of cocaine-seeking behaviour, even though shorter sessions achieve less drug intake overall (Allain & Samaha, 2019). This suggests that intermittent cocaine use—even in brief daily bouts permitting only limited cocaine intake—is sufficient to facilitate the development of addiction-like behaviours. However, it remains unclear how long versus short bouts of intermittent cocaine intake affect cue-triggered cocaine-seeking behaviour. Addressing this question could provide new information on how different histories of drug exposure might influence cue-triggered cravings and relapse, potentially informing personalized interventions. If prolonged cocaine self-administration is required to produce the brain and psychological changes that underlie cue-triggered relapse, then Long-IntA sessions should promote greater cue-induced cocaine-seeking behaviour compared to Short-IntA sessions. To test this hypothesis, we measured cocaine seeking triggered by discriminative stimuli (DS) previously signalling when instrumental responding will be reinforced with cocaine and conditioned stimuli (CS) previously signalling drug delivery.

## Methodology

### Animals

All experimental procedures were approved by the animal ethics committee of the Université de Montréal and carried out in accordance with the recommendations of the Canadian Council on Animal Care. Female (150-175 g; *n* = 38) Sprague Dawley rats (Envigo, Frederick Barrier 208A, MD, USA) were individually housed on a reversed 12-hour light/dark cycle (8:30 AM lights off) and held in a climate-controlled colony. Rats were trained and tested during the dark phase of the light/dark cycle. Upon their arrival, rats had a 3-day acclimation period where they had unlimited access to standard laboratory chow. Food was then restricted to 15 g/day for all experimental phases (Algallal et al., 2020; Martin et al., 2010). Water was available ad libitum.

### Apparatus

All behavioural training and testing took place in standard operant conditioning chambers (Med Associates, St Albans, VT, USA) placed in sound attenuating boxes equipped with a ventilation fan. The front wall of each chamber had two retractable levers with a white cue light above each. The back wall also had two white cue lights, one at the top centre and another on the bottom right. The chambers were also equipped with a clicker and a tone generator (2900 Hz, 75 dB). During sucrose self-administration training, an electronic candle located outside of the cage was illuminated for the duration of the session. A pellet dispenser distributed sucrose pellets (45 mg, banana flavoured; Bioserv; product # 76285-260, VWR, Mount Royal, QC, Canada) into a recessed magazine. Infrared cells located inside the magazine port counted the number of magazine entries and four rows of infrared cells at the bottom of each cage measured locomotion.

### Magazine Training

Fig. 1A shows the sequence of experimental events. To prepare rats for subsequent autoshaping sessions where sucrose would be used as an unconditioned stimulus (US), rats were first trained to retrieve sucrose from the magazine. Independent of the rats’ behaviour, pellets were delivered into the magazine on a variable interval 45-seconds (VI45-s) schedule until a total of 40 pellets were delivered per session (2 sessions total).

**Figure 1.**
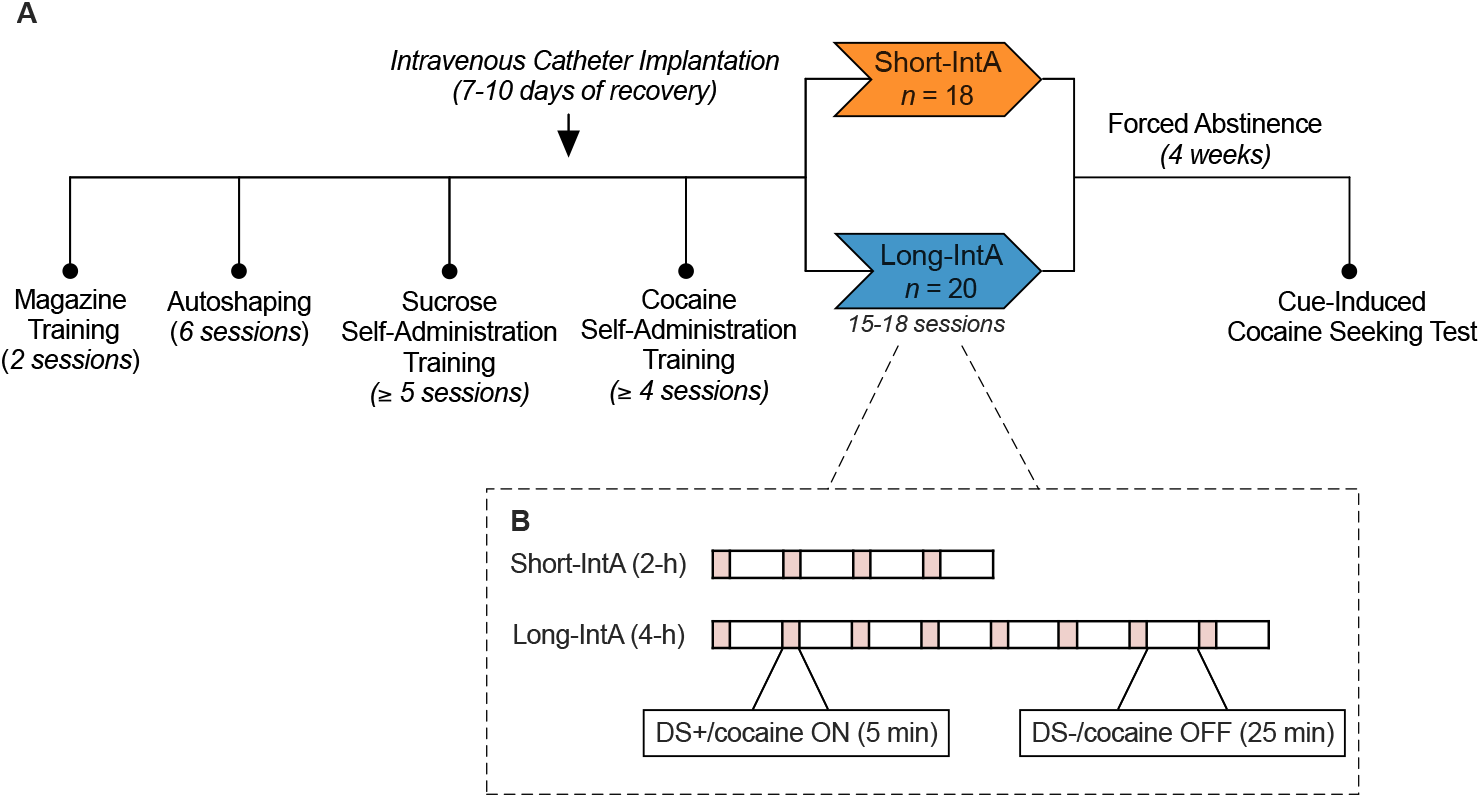
The sequence of experimental events. **(A)** Overview of the experimental timeline. **(B)** Sequence of events within a single intermittent access (IntA) cocaine self-administration session under Short-IntA and Long-IntA conditions. IntA, Intermittent access; DS+, discriminative stimulus signalling cocaine availability; DS-, discriminative stimulus signalling cocaine unavailability.

### Autoshaping

When a conditioned stimulus (CS) predicts a US, rats can show different conditioned approach responses; sign trackers preferentially approach and interact with the CS, whereas goal trackers preferentially approach the site of reward delivery, and intermediates approach both. In appetitive settings, sign-trackers are thought to respond more to CSs and goal-trackers are thought to respond more to DSs (Flagel et al., 2011; Pitchers et al., 2017; c.f. Kawa et al., 2016; LeCocq et al., 2024, 2025; Martin et al., 2022).

To consider this potential individual variability, rats received six, 30-min autoshaping sessions to determine their Pavlovian conditioned approach phenotype. During these sessions, extension of a lever CS for 8 s was immediately followed by the delivery of a sucrose pellet (US) on a VI 45-s schedule, for a total of 30 reinforcers/session. The lever CS (left- or right-side lever) was assigned as the active lever in subsequent experimental phases. Using data averaged over the last 2 autoshaping sessions, a response bias score was calculated for each rat [(CS lever activations – CS port entries) / (CS lever activations + CS port entries)]. Sign-trackers had a score > +0.50, goal-trackers, a score < -0.50, and intermediates, a score between -0.49 to +0.49.

### Sucrose Self-Administration Training

To facilitate the subsequent acquisition of lever-pressing behaviour to self-administer cocaine, rats were trained to press on the active lever to receive one sucrose pellet into the magazine during 30-min sessions. For the first two sessions, or until rats met acquisition criteria, pellets were available on a fixed ratio 1 (FR1) schedule of reinforcement. Acquisition criteria were administration of 20 pellets/session and twice as many active vs. inactive lever presses, for two consecutive sessions. Once criteria were met, the reinforcement schedule was increased to FR3, until the same acquisition criteria were met again.

### Surgical Intravenous Catheter Implantation

Following the acquisition of sucrose self-administration behaviour, the rats were placed under isoflurane anesthesia (5% for induction, 2% for maintenance) and homemade silastic catheters were implanted into the right jugular vein (See Samaha et al., 2011; Weeks, 1962 for detailed procedures). Before the start of surgery, rats received an intramuscular injection of penicillin (Procollin; 0.02 ml of a 300 mg/ml solution; CDMV, Saint-Hyacinthe, QC, Canada) and a subcutaneous injection of Carprofen (Rimadyl; 0.03 ml of a 50 mg/ml solution; CDMV). Rats received 7-10 days of recovery. To ensure catheter patency, catheters were flushed daily from implantation onward, alternating between a solution of 0.05 ml heparin with Baytril (heparin: 0.1 ml of a 0.2 mg/ml solution; Sigma-Aldrich, Oakville, ON; Baytril: 2 mg/ml; CDMV) and 0.05 ml saline.

### Drugs

Cocaine (Galenova Inc., St Laurent, QC) was dissolved in 0.9% sterile saline and filtered using a corning bottle-top filter (0.22-μm PES membrane; Fisher Scientific, Whitby, ON, Canada). The concentration of the cocaine solution was adjusted every third day based on average rat body weight.

### Cocaine Self-Administration Training

Following recovery from surgery, the rats were trained to self-administer intravenous cocaine during daily 1-h sessions. The rats’ catheters were connected to an infusion pump via Tygon tubing that was protected by a metal spring and a liquid swivel to allow free movement. At the start of each session, the fan was activated. Two minutes later, both levers extended and the cue light above the left lever illuminated to serve as a DS+ signalling cocaine availability and remained illuminated for the entire session. Because some rats had the left lever designated as active and others had the right lever as active, the DS+ was counterbalanced above the active and inactive levers across rats. Cocaine (0.5 mg/kg/infusion over 5 seconds, in a volume of ∼150 µl) was available under an FR3 schedule of reinforcement, and each infusion was paired with a 5-s CS+. CS+ modality was counterbalanced across rats, consisting of either illumination of the cue light above the right lever combined with a tone or illumination of the cue light on the bottom right of the back wall combined with a clicker. The criteria to proceed to intermittent access (IntA) self-administration training were i) obtaining ≥ 6 cocaine infusions/session, ii) taking cocaine regularly across the session, as indicated by cumulative response records, and iii) making twice as many active vs. inactive lever presses.

### Intermittent-Access (*IntA*) Cocaine Self-Administration

Following cocaine self-administration training, rats were divided into 2 groups and received either 2-h (Short-IntA; *n* = 18) or 4-h (Long-IntA; *n* = 20) IntA sessions during which DS+/cocaine ON periods alternated with DS-/cocaine OFF periods (see Fig. 1B). At the start of each session, the fan was activated. Two minutes later, both levers extended along with illumination of the same DS+ used during the previous cocaine self-administration sessions. The DS+ stayed on for 5 minutes, during which pressing the active lever produced a cocaine infusion (0.5 mg/kg/infusion over 5 s) under an FR3 schedule of reinforcement. Each cocaine infusion was paired with a 5-s presentation of the same CS+ presented during the previous cocaine self-administration sessions. The DS- (the cue light at the top centre of the back wall) was illuminated. The DS-stayed on for 25 minutes, during which pressing the active lever produced a 5-s CS-under an FR3 schedule of reinforcement and no cocaine. CS-modality was counterbalanced with CS+ modality, such that for half the rats it consisted of illumination of the cue light above the right lever combined with a tone, and for the other half it consisted of illumination of the cue light on the bottom right of the back wall combined with a clicker. This cycle of 5 minutes DS+/cocaine ON and 25 minutes DS-/cocaine OFF was repeated 4 times in the Short-IntA group and 8 times in the Long-IntA group.

Cocaine self-administration was considered to have come under the control of the DSs when rats made ≥ 70% of their active lever presses during the DS+ period and took at least one cocaine injection per DS+ period. Rats received 16-18 training sessions. One rat completed only 15 sessions due to loss of catheter patency. As all rats reached the learning criteria by Session 15, self-administration data were analyzed for the first 15 sessions across all animals. A computer issue caused the loss of all data from Session 13 so data from this session is not available.

### Catheter Patency

At the end of IntA training, catheter patency was verified. Catheters were flushed with the short-acting barbiturate propofol (0.05 ml of a 10 mg/ml solution, Fresenius Kabi Canada Ltd, Richmond Hill, ON) followed by 0.05 ml saline. If ataxia was observed within 5-10 seconds following the injection, the rats were considered to have patent catheters. If rats failed to show ataxia, they were immediately given the same dose a 2^nd^ time. Rats that failed to show ataxia after the 2^nd^ dose were excluded from the study.

### Cue-Induced Cocaine-Seeking Test

Following the last IntA session, rats stayed in their home cages without access to cocaine. Following 4 weeks of forced abstinence, rats received a cue-induced cocaine-seeking test. Immediately before the start of the test, rats received an injection of cocaine (10 mg/kg, i.p.; Kalivas & McFarland, 2003), such that they experienced the cues with drug onboard, as during IntA cocaine self-administration. The test began with the fan turning on. After 2 min, both levers extended, and a cue (i.e., DS+, CS+, DS-, CS- or DS+CS+) was presented response-independently for 2 min. Cue presentation was followed by a 1-minute inter-trial interval (ITI), during which the levers remained extended, but no cues were presented. Each cue was presented 3 times, in counterbalanced order. Lever presses produced no cocaine or cues.

### Statistical Analyses

We used Shapiro-Wilk tests to determine the normality of data distribution. We analyzed normally distributed data using independent samples two-tailed *t*-tests. We analyzed non-normally distributed data using Wilcoxon signed-rank tests and Mann-Whitney U tests. We corrected for multiple comparisons using the Holm-Bonferroni method. We set the statistical significance threshold as *p* ≤ 0.05 and analyzed data using IBM SPSS (Version 29). Increased variability observed during the cue-induced cocaine-seeking test prompted two separate quartile analyses based on the number of active lever presses during (1) DS+ induced cocaine-seeking and (2) CS+ induced cocaine-seeking, to identify subgroups of rats with differing relapse propensities. In each case, we classified rats into three subgroups reflecting their rate of cocaine-seeking behaviour on the relapse test: High relapse (active lever presses ≥ third quartile), low relapse (≤ the first quartile), and intermediate (strictly between the first and third quartiles). We applied linear support vector machine binary classification, implemented using the LinearSVC class from the scikit-learn package in Python, to try to predict high and low (DS+) and (CS+)-induced relapse based on behaviour during the 15^th^ IntA session [i.e., hourly rates of cocaine intake, active lever pressing during DS+ and DS− periods and locomotion, as well as average latency to obtain the first cocaine infusion upon DS+ onset, average number of burst-like episodes of consumption across DS+ periods (taking › 5 infusions in ≤ 5 minutes; Belin et al., 2009) and discrimination ratio. This classification approach computes, for each rat and for each cue type (i.e., DS+ and CS+), low and high relapse prediction scores that are linear combinations of the behavioural measures. We used scatter plots to visualize the relationship between relapse prediction scores and actual relapse behaviours, as measured by the number of active lever presses. We used receiver operating characteristic curves to evaluate the performance of the relapse prediction model in relation to the classification thresholds.

## Results

### Autoshaping

Prior to any self-administration experience, rats received 6 autoshaping sessions where a lever CS predicted sucrose reward. Fig. 2A shows average response bias scores for sign-tracker, goal-tracker, and intermediate rats. By the last autoshaping session, most rats were sign-trackers (71%), followed by goal-trackers (16%) and intermediates (13%). Across sessions, sign-trackers increased their number of lever presses (Fig. 2B; Session 1 vs. 6: *z* = 3.63, *p* < 0.001) while goal-trackers and intermediates maintained stable responding (Fig. 2B; Session 1 vs. 6: INT *z* = 0.67, *p* = 0.50; GT *z* = -1.76, *p* = 0.08). On Session 6, sign-trackers showed more lever activations than did the other phenotypes (Fig. 2B; ST vs GT: *U* = 1.5, *p* < 0.001; ST vs INT: *U* = 21, *p* = 0.01), and intermediates also had more lever activations compared to goal-trackers (Fig. 2B; GT vs. INT: *U* = 1, *p* = 0.01). Both sign-trackers and intermediates decreased their number of magazine entries over sessions (Fig. 2C; Session 1 vs. 6: ST *z* = 0, *p* < 0.001; INT *z* = 0, *p* < 0.04; GT *z* = 16, *p* = 0.25). Consequently, on Session 6, magazine entries were highest in goal-trackers and lowest in sign-trackers (Fig. 2C; GT vs. ST: *U* = 1, *p* < 0.001; GT vs INT: *U* = 3, *p* < 0.02; ST vs INT: *U* = 5.5, *p* < 0.001). Thus, all rats learned a Pavlovian conditioned approach response with CS-US conditioning, but the nature of that response varied, with some rats learning to approach the CS and others learning to approach the site of US delivery.

**Figure 2.**
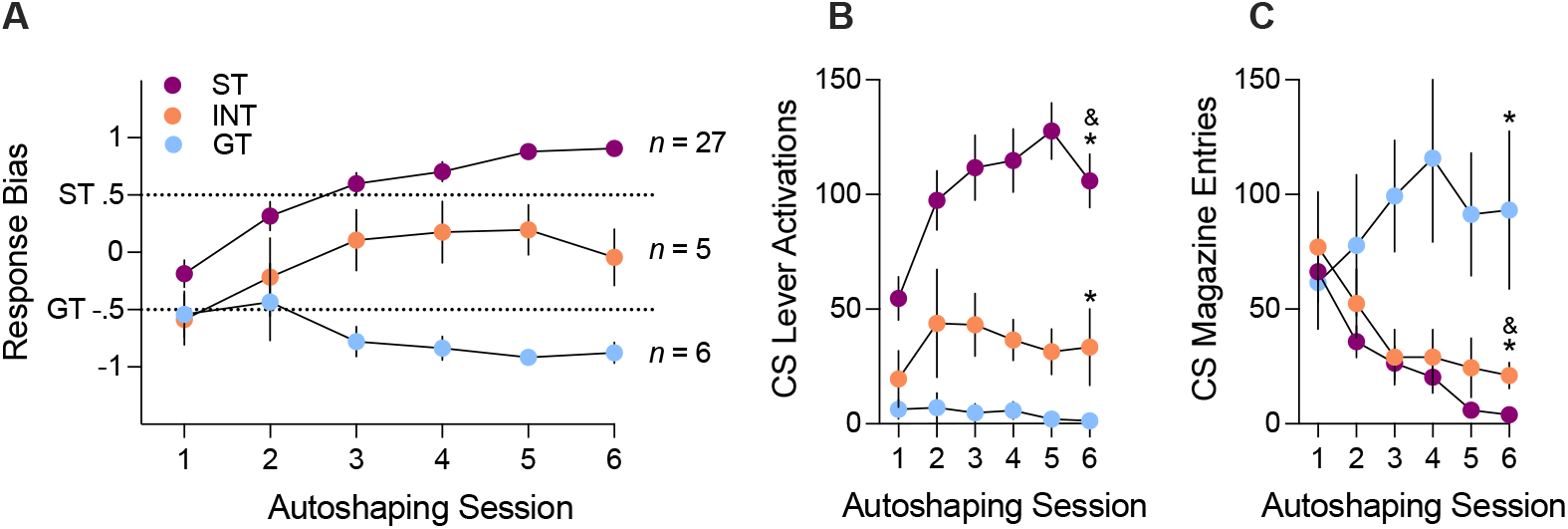
Acquisition of Pavlovian conditioned approach behaviour. **(A)** Average (± SEM) conditioned approach response bias for sign-trackers (ST; purple), goal-trackers (GT; blue) and intermediates (INT; orange). The dashed lines represent cut-off values for each phenotype. **(B)** Average (± SEM) lever activations and **(C)** magazine entries for each phenotype made during presentation of the lever conditioned stimulus (CS). & vs. Session 1 in the same group; * vs. other phenotypes on Session 6; *N* = 38.

### Cocaine Self-Administration Behaviour in Short-vs. Long-IntA Groups

On Session 1, Short-IntA (Fig. 3A) and Long-IntA (Fig. 3B) rats made approximately half of their active lever presses during DS+ periods (Fig. 3A inset, Short-IntA 45%; Fig. 3B inset, Long-IntA 42%) and the other half during DS-periods, even though cocaine was only available during DS+ periods. However, on Session 15, both Short-IntA rats (Fig. 3C) and Long-IntA rats (Fig. 3D) responded most during DS+ periods (Fig. 3C inset, Short-IntA 76%; Fig. 3D inset; Long-IntA 72%) and inhibited responding during DS-periods. In accordance with this, on Session 1, Short-IntA rats showed no significant difference in discrimination ratios for active lever pressing during the DS+ vs. DS- (Fig. 3E; *z* = 1.46, *p* = 0.07), and Long-IntA rats had a significantly higher DS-vs. DS+ ratio (Fig. 3F; *z* = 2.05, *p* = 0.02). This indicates that in both groups, the DSs did not yet guide cocaine self-administration behaviour. However, on Session 15, the DS+ discrimination ratio was significantly greater than it was on Session 1 in both Short-IntA (Fig. 3E; *z* = 2.72, *p* = 0.01) and Long-IntA (Fig. 3F; *z* = 3.40, *p* < 0.001) rats, and it was also greater than the DS-ratio (Fig. 3E, Short-IntA; *z* = -2.90, *p* = 0.002; Fig. 3F, Long-IntA; *z* = -3.14, *p* = 0.001). In parallel, the DS-ratio decreased across sessions in Short-IntA (Fig. 3E; Session 1 vs. 15: *z* = -2.72, *p* = 0.003) and Long-IntA (Fig. 3F; Session 1 vs. 15, *z* = - 3.40, *p* < 0.001) rats. Thus, cocaine self-administration behaviour came under the control of the DSs in both Short-IntA and Long-IntA rats.

**Figure 3.**
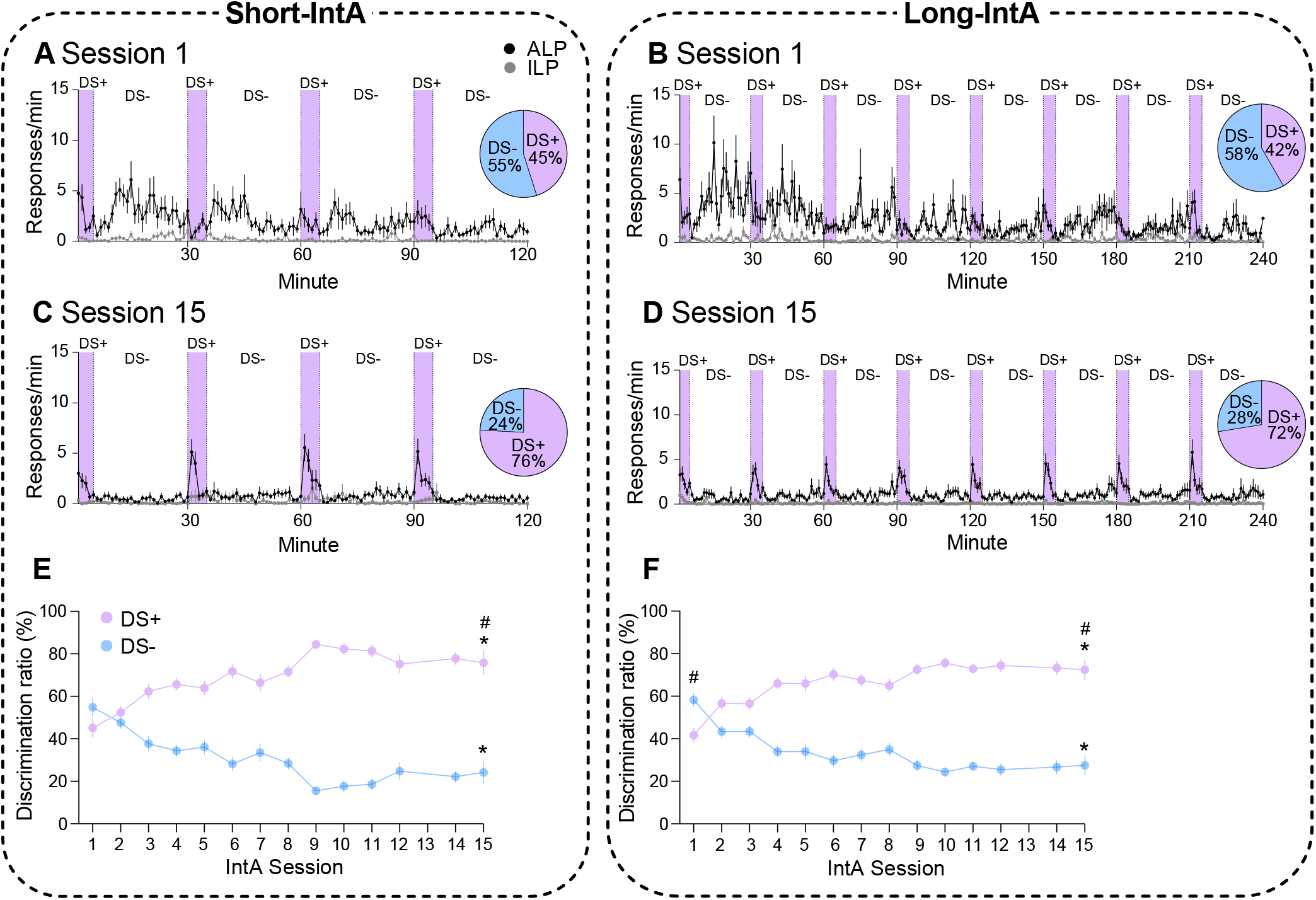
Cocaine self-administration comes under the control of discriminative stimuli signalling drug availability (DS+) and unavailability (DS-). Average (± SEM) lever presses made during DS+ (purple-shaded) and DS- (blue-shaded) periods on Intermittent-Access (IntA) Session 1 in **(A)** Short-IntA and **(B)** Long-IntA rats, and on IntA Session 15 in **(C)** Short-IntA and **(D)** Long-IntA rats. Pie chart insets represent the percentage of active lever presses made during the DS+ (purple) vs. DS- (blue) periods. Average (± SEM) DS+ (purple) and DS- (blue) discrimination ratios across sessions in **(E)** Short-IntA rats and **(F)** Long-IntA rats. *vs. Session 1 of the same condition; #DS+ vs. DS-on the same session; *n* = 18 Short-IntA, *n* = 20 Long-IntA.

Compared to Short-IntA rats, Long-IntA rats self-administered more cocaine injections during both the first and 15^th^ IntA sessions (Fig. 4A; Session 1, *U* = 111, *p* = 0.02; Session 15, *U* = 91.5, *p* = 0.01). Long-IntA rats also escalated their cocaine intake from Session 1 to 15 (Fig. 4A; *z* = 1.97, *p* = 0.049), whereas Short-IntA rats maintained stable levels of intake (Fig. 4A; *z* = 0.65, *p* = 0.52). Accordingly, average cumulative cocaine intake was greatest in the Long-IntA rats, with these rats taking over twice as much drug as did Short-IntA rats (Fig. 4B; *t*_(36)_ = - 4.16, *p* < 0.001). Thus, compared to Short-IntA rats, Long-IntA rats consumed significantly more cocaine across IntA sessions.

**Figure 4.**
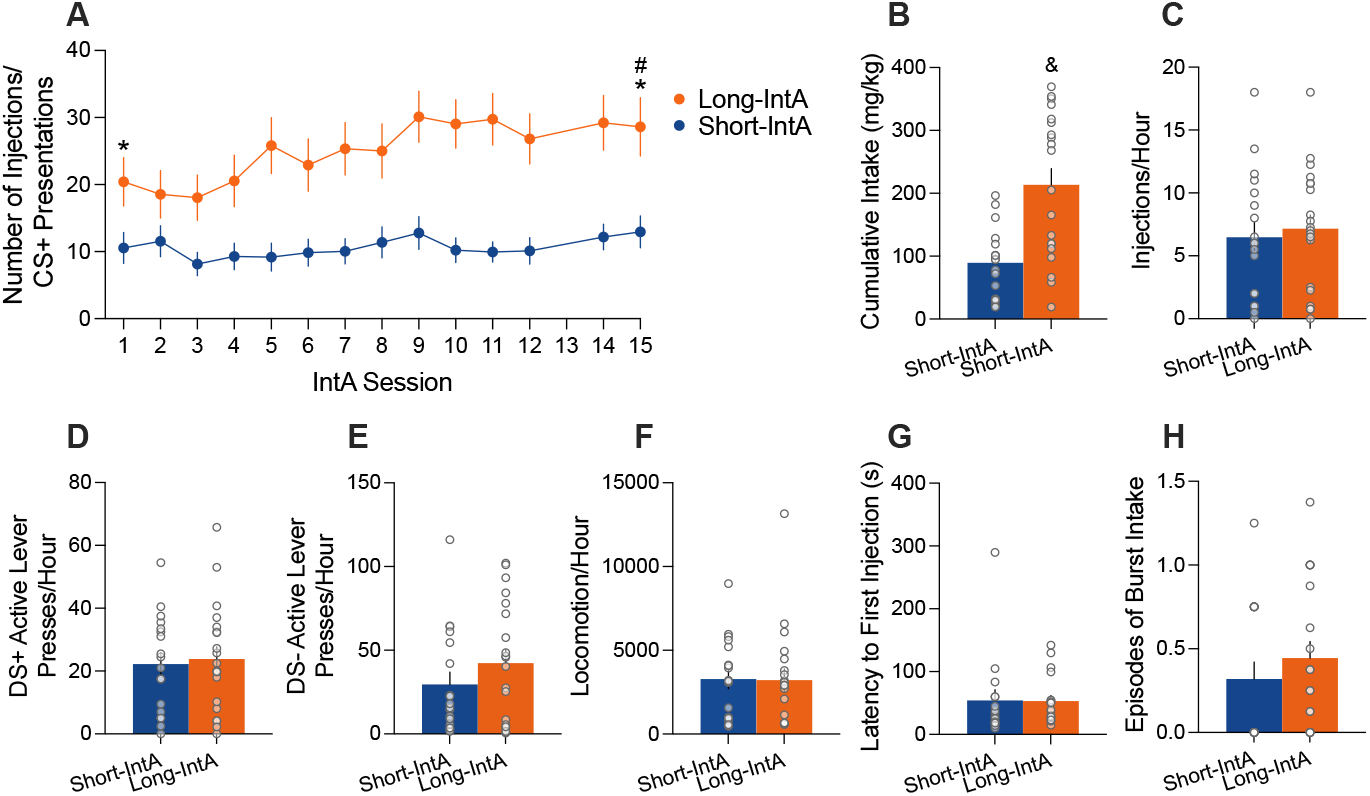
More cumulative cocaine intake in Long-IntA rats but similar hourly rates of responding in Long- and Short-IntA groups. **(A)** Average (±SEM) injections/CS+ presentations self-administered across Intermittent-Access (IntA) sessions in Short-IntA (blue circles) and Long-IntA (orange circles) rats. **(B)** Average (± SEM) cumulative cocaine intake (mg/kg) in Short-IntA (blue bars) and Long-IntA (orange bars) rats. Average (± SEM) **(C)** hourly rates of self-administered cocaine injections, hourly rates of active lever presses made during **(D)** DS+ periods and **(E)** DS- periods, **(F)** hourly rates of locomotion, **(G)** average (± SEM) latency to obtain the first injection upon DS+ onset and **(H)** number of burst-like episodes of cocaine intake averaged (± SEM) across DS+ periods, in Short-IntA (blue bars) and Long-IntA (orange bars) rats. Circles in bars graphs represent data from individual rats. * Long-IntA > Short-IntA on the same IntA session. #Session 15 vs. Session 1 in Long-IntA rats. & Long-IntA > Short-IntA. CS+, conditioned stimulus paired with cocaine delivery; DS+ discriminative stimulus predicting cocaine availability; DS- discriminative stimulus predicting cocaine unavailability; *n* = 18 Short-IntA, *n* = 20 Long-IntA.

On the 15^th^ IntA session, we also analyzed group differences in hourly rates of cocaine intake (Fig. 4C), hourly rates of active lever presses during DS+ (Fig. 4D) and DS- (Fig. 4E) periods, hourly rates of locomotion (Fig. 4F), average latency to obtain the first injection upon DS+ onset (Fig. 4G) and average number of episodes of burst-like cocaine intake across DS+ periods (Fig. 4H). There were no group differences on any of these measures (Fig. 4C, *t*_(36)_ = -0.42, *p* = 0.68; Fig. 4D, *t*_(36)_ = -0.31, *p* = 0.76; Fig. 4E, *U* = 151.5, *p* = 0.41; Fig. 4F, *U* = 167, *p* = 0.72; Fig. 4G, *U* = 121, *p* = 0.32; Fig. 4H, *U* = 136, *p* = 0.21). This suggests that group differences in cumulative cocaine intake primarily reflected differences in session length rather than distinct self-administration profiles between groups.

Finally, to further examine the temporal pattern of cocaine intake in the two groups, we analyzed minute-by-minute cocaine self-administration during the DS+/cocaine ON periods. Fig. 5 shows the average number of infusions earned during each min of the 5-min DS+/cocaine ON periods in Sessions 2, 6, and 15. Rats in both groups took most of their cocaine infusions in the first min, and this loading effect increased across sessions. On Session 2, Short-IntA rats (Fig. 5A) took more cocaine during Min 1 compared to Min 4 (*z* = -3.14, *p* = 0.002) and Min 5 (*z* = - 2.74, *p* = 0.01). On Session 6, they took more drug during Min 1 compared to Min 3 (*z* = -3.33, *p* < 0.001), Min 4 (*z* = - 3.30, *p* < 0.001) and Min 5 (*z* = -4.08, *p* < 0.001). On Session 15, Short-IntA rats took more cocaine during Min 1 compared to each subsequent min (vs. Min 2: *z* = -2.36, *p* = 0.02; vs. Min 3: *z* = -4.12, *p* < 0.001; vs. Min 4: *z* = -4.41, *p* < 0.001; vs. Min 5: *z* = -4.66, *p* < 0.001). Finally, from Sessions 2 to 15, rats escalated their cocaine intake during the first min (*z* = 2.70, *p* = 0.004).

**Figure 5.**
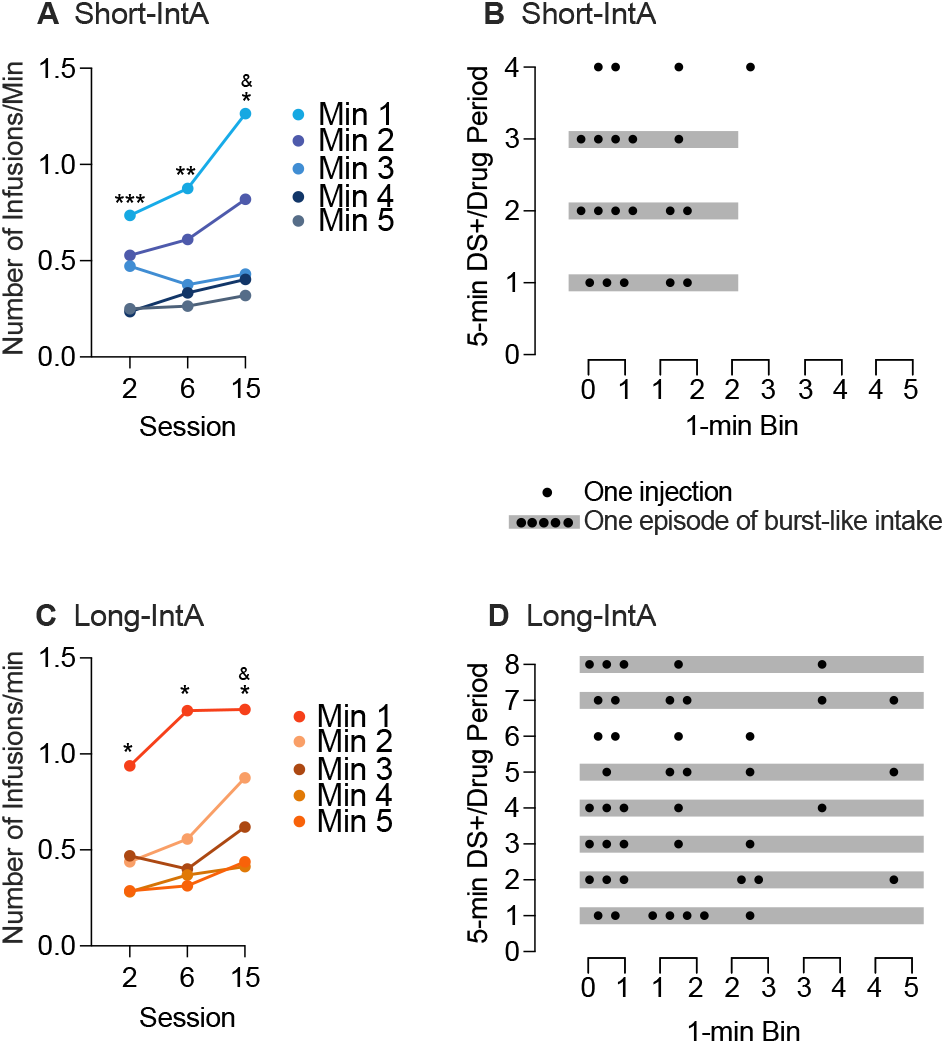
Short-IntA and Long-IntA rats take most of their cocaine infusions in the first minute of the 5-min DS+/cocaine ON periods. Average (± SEM) infusions self-administered by the **(A)** Short-IntA and **(C)** Long-IntA rats during Minutes 1-5 of the 5-min DS+ periods on Session 2, 6, and 15. Episodes of burst-like intake (taking > 5 injections in ≤ 5 min) during the 15^th^ IntA session in a representative **(B)** Short-IntA and **(D)** Long-IntA rat. ***min 1 vs. min 4-5 on the same session; **min 1 vs. min 3-5 on the same session; *min 1 vs. all other minutes on the same session; &Session 2 vs. 15. IntA, Intermittent Access. *n* = 18 Short-IntA, *n =* 20 Long-IntA.

The Long-IntA rats (Fig. 5C) showed a similar pattern of cocaine self-administration. On Session 2, they took more cocaine during Min 1 than in each subsequent min (vs. Min 2: *z* = -5.11, *p* < 0.001; vs. Min 3: *z* = -4.55, *p* < 0.001; vs. Min 4:*z* = -6.59, *p* < 0.001; vs. Min 5: *z* = -6.27, *p* < 0.001). The Long-IntA rats maintained this cocaine loading pattern both on Session 6 (Min 1 vs. 2: *z* = -5.67, *p* < 0.001; Min 1 vs. 3: *z* = -6.80, *p* < 0.001; Min 1 vs. 4: *z* = -7.08, *p* < 0.001; Min 1 vs. 5: *z* = -7.28, *p* < 0.001) and Session 15 (Min 1.vs 2: *z* = -3.30, *p* < 0.001; Min 1 vs. 3: *z* = -5.17, *p* < 0.001; Min 1 vs. 4: *z* = -6.67, *p* < 0.001 ; Min 1 vs. 5: *z* = -6.66, *p* < 0.001). Across sessions, they also escalated their cocaine intake during the first min (Session 2 vs. 15: *z* = 2.61, *p* = 0.005).

In summary, both Short-IntA and Long-IntA rats learned to load up on cocaine during the 1^st^ minute of signalled drug availability, maximizing their intake as soon as cocaine became available, and this effect sensitized across sessions (Allain et al., 2018; Allain & Samaha, 2019; D’Ottavio et al., 2023). Thus, while Long-IntA rats had twice the amount of time to self-administer cocaine compared to Short-IntA rats (i.e., 8 x 5-min cocaine ON periods/session vs. 4), the two groups showed a similar temporal pattern of cocaine intake.

### Cue-Induced Cocaine-Seeking Test

After 4 weeks of forced abstinence from the drug, we injected the rats with cocaine and assessed group differences in drug-seeking behaviour following response-independent presentation of the DS+, DS-, CS+, CS- and DS+CS+ combined. First, we found no significant correlations between Pavlovian conditioned response score and active lever pressing during cue presentation on the cocaine-seeking test (Table 1). This is in accordance with prior work showing that a goal-tracker vs. sign-tracker phenotype does not predict reward-seeking behaviour upon DS or CS presentation (Algallal et al., 2025; Flagel et al., 2011; Kawa et al., 2016; LeCocq et al., 2024, 2025; Martin et al., 2022; Pitchers et al., 2017).

**Table 1.** Spearman Correlations Between Pavlovian Conditioned Response Bias and Lever-Pressing Behaviour During a Cue-Induced Cocaine Seeking Test.

|  | DS+ | CS+ | DS- | CS- | DS+CS+ |
| --- | --- | --- | --- | --- | --- |
| Short-IntA | $r = -0.07$<br>$p = 0.79$ | $r = 0.21$<br>$p = 0.39$ | $r = 0.03$<br>$p = 0.90$ | $r = 0.44$<br>$p = 0.07$ | $r = 0.12$<br>$p = 0.64$ |
| Long-IntA | $r = -0.39$<br>$p = 0.09$ | $r = -0.27$<br>$p = 0.26$ | $r = -0.22$<br>$p = 0.36$ | $r = 0.05$<br>$p = 0.82$ | $r = -0.31$<br>$p = 0.18$ |

Second, in Short-IntA rats (Fig. 6A), DS+ presentation triggered the greatest increases in active lever pressing (vs. ITI, *z* = -2.61, *p* = 0.01; vs. CS+, *z* = -2.18, *p* = 0.03; vs. DS-, *z* = -3.08, *p* = 0.003; vs. CS+DS+, *z* = -2.17, *p* = 0.02). Rats pressed more during CS+ vs. CS-(*z* = -2.56, *p* = 0.01) or DS- (*z* = -2.04, *p* = 0.02), but responding during the CS+ and DS+CS+ was similar to that during the inter-trial interval, when no cue was presented (ITI vs. CS+, *z* = - 0.74, *p* = 0.23; ITI vs. DS+CS+, *z* = 0.23, *p* =.41). The DS- and CS-inhibited cocaine seeking relative to the inter-trial interval (DS-vs. ITI, *z* = -2.56, *p* = 0.01; CS-vs. ITI, *z* = - 3.31, *p* < 0.001). Thus, when presented response-independently, both the CS- and DS-functioned as effective conditioned inhibitors of cocaine seeking, whereas the DS+, but not CS+, served as a conditioned excitor, prompting increases in drug-seeking responses.

**Figure 6.**
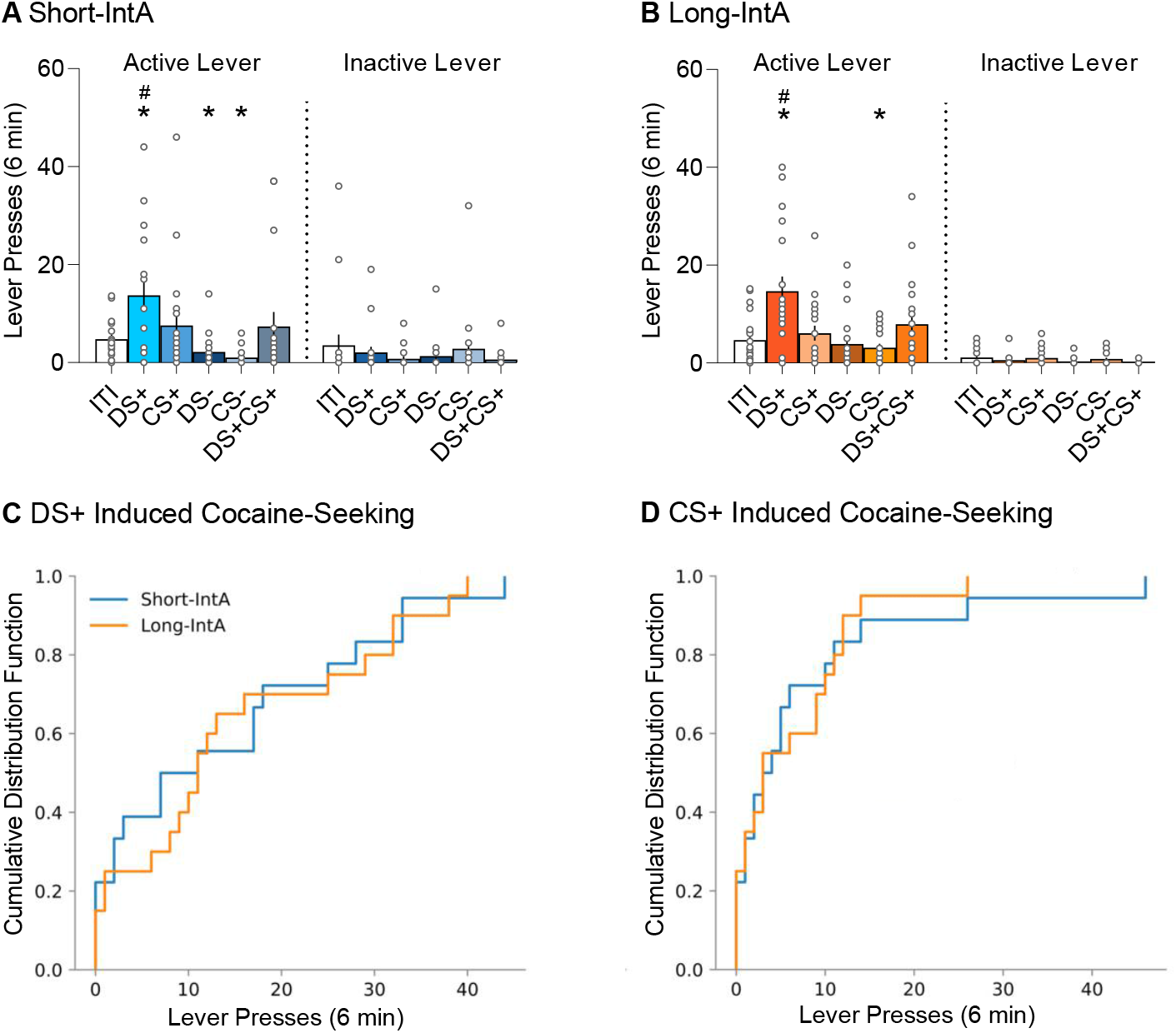
Response-independent presentation of a cocaine-associated DS+ (but not CS+) triggers increases in drug-seeking behaviour and responding is similar in Short-IntA and Long-IntA rats. Average (± SEM) active and inactive lever presses during inter-trial intervals (ITI), and DS+, CS+, DS-, CS-, and DS+CS+ presentation in **(A)** Short-IntA rats and **(B)** Long-IntA rats. Cumulative distribution functions of active lever pressing during **(C)** DS+ and **(D)** CS+ presentation in Short-IntA rats (blue line) and Long-IntA rats (orange line). *vs. ITI; #vs. CS+. IntA, Intermittent Access; ALP, active lever presses; ILP, inactive lever presses; DS+, discriminative stimulus signalling cocaine availability; DS-, discriminative stimulus signalling cocaine unavailability; CS+, conditioned stimulus paired with cocaine delivery; CS-, conditioned stimulus never paired with cocaine. *n* = 18 Short-IntA, *n* = 20 Long-IntA.

Similarly, in Long-IntA rats (Fig. 6B), the DS+ triggered significantly more cocaine-seeking behaviour compared to the other cue conditions (vs. ITI, *z* = -3.24, *p* = 0.001; vs. CS+, *z* = -2.70, *p* = 0.01; vs. DS-, *z* = -3.30, *p* < 0.001; vs. CS+DS+, *z* = -2.64, *p* = 0.004). During CS+ presentation, responding was greater than during CS- (*z* = -1.76, *p* = 0.04) but not DS-presentation (*z* = -1.53, *p* = 0.06). However, neither the CS+ (*z* = -1.23, *p* = 0.11) nor the DS+CS+ combined (*z* = 1.45, *p* = 0.07) increased lever-pressing behaviour compared to the inter-trial interval. The CS-(but not DS-) inhibited cocaine seeking relative to the inter-trial interval (CS-vs. ITI, *z* = -1.67, *p* = 0.05; DS-vs. ITI, *z* = -1.29, *p* = 0.10). Thus, in both Short-IntA and Long-IntA groups, DS+ presentation was more effective in invigorating the pursuit of cocaine compared to CS+ presentation.

The two groups responded similarly during the cue-induced cocaine-seeking test (Fig. 6A vs. B; DS+, *U* = 189.5, *p* = 0.39; CS+, *U* = 181, *p* = 0.5; DS+CS+ combined, *U* = 212.5, *p* = 0.17, and ITI, *U* = 158.5, *p* = 0.27), except that the Long-IntA rats responded more to the CS-than did the Short-IntA rats (*U* = 256.5, *p* = 0.01). Thus, although Long-IntA rats took significantly more cocaine in the past than did Short-IntA rats, and also had more cue-cocaine pairings, the two groups showed similar rates of cue-induced cocaine-seeking behaviour.

Cumulative distribution functions for active lever responses during DS+ (Fig. 6C) and CS+ (Fig. 6D) further highlight the similarity of cocaine-seeking behaviour in the Short-IntA and Long-IntA rats. As indicated by the overlap between the curves representing each group, Short-IntA and Long-IntA rats showed comparable distributions of active lever pressing in the presence of the DS+ and CS+. The cumulative distribution functions also point to the higher rates of responding during DS+ vs. CS+ across groups. For example, 39% of Short- and Long-IntA rats made ≥ 15 active lever presses/6 min during DS+ presentation but only 8% did so during CS+ presentation. Thus, despite producing significant differences in cumulative cocaine intake, Short IntA and Long-IntA cocaine self-administration produce comparable interindividual variability in later responding to cocaine cues.

### Behavioural Markers During IntA Predict the Risk to Relapse

We analyzed the extent to which behaviour during IntA cocaine self-administration predicted cue-induced cocaine seeking. Because Short-IntA and Long-IntA rats showed no differences in responding on the cue-induced cocaine-seeking test, we pooled data across groups. First, we stratified the number of active lever presses/6min during DS+ and CS+ presentation into quartiles (Quartile 1; ‘Low relapse’, Quartiles 2-3; pooled into ‘Intermediate relapse’, Quartile 4; ‘High relapse’). We then examined if behavioural measures during the 15^th^ IntA session predicted quartile membership. These measures were hourly rates of cocaine intake, hourly rates of active lever pressing during DS+ and DS-periods, hourly rates of locomotion, as well as the average latency to make the first injection upon DS+ onset, the average number of burst-like episodes of intake across DS+ periods and DS+ discrimination ratio. This revealed distinct phenotypes that predicted the rate of (DS+)-induced relapse-like behaviour. First, rats that had self-administered more drug infusions/hour later showed the greatest (DS+)-induced increases in cocaine-seeking behaviour (Fig. 7A; Intermediates vs. low-relapsing rats, *t*_(25)_ = -1.95, *p* = 0.03; high-relapsing vs. low-relapsing rats, *t*_(20)_ = -3.36, *p* = 0.003; Intermediates vs. high-relapsing rats,(*t*_(25)_ = -1.63, *p* = 0.12). Second, rats with high rates of active lever pressing during DS+ trials later responded more to the DS+ on the cocaine-seeking test (Fig. 7B; Intermediates vs. low-relapsing rats, *t*_(25)_ = -2.13, *p* = 0.02; high-relapsing rats vs. low-relapsing rats, *t*_(20)_ = = -3.45, *p* = 0.003; high-relapsing rats vs. intermediates, *t*_(25)_ = -1.75, *p* = 0.046). Third, rats with high rates of active lever pressing during DS-trials also later responded more to the DS+ on the cocaine-seeking test (Fig. 7C; Intermediates vs. low-relapsing rats, *U* = 42, *p* = 0.02; high-relapsing rats vs. low-relapsing rats, *U* = 9, *p* < 0.001; high-relapsing rats vs. intermediates, *U* = 48, *p* = 0.03). Fourth, rats with high prior rates of cocaine-induced locomotion showed greater responding to the DS+ on the cocaine-seeking test (Fig. 7D; high-relapsing rats vs. low-relapsing rats; *U* = 29, *p* = 0.02; no other comparisons were statistically significant). Fifth, rats that self-administered their first cocaine injection more rapidly upon DS+ onset later responded more to the DS+ on the cocaine-seeking test (Fig. 7E; intermediates vs. low-relapsing rats, *U =* 52, *p* = 0.06; high-relapsing rats vs. low-relapsing rats, *U* = 27, *p* < 0.01; intermediates vs. high-relapsing rats, *U* = 70, *p* = 0.27). Sixth, more episodes of burst-like cocaine intake averaged across DS+ periods predicted greater rates of responding to the DS+ on the cocaine-seeking test (Fig. 7F; intermediates vs. low-relapsing rats, *U =* 63, *p* = 0.12; high-relapsing rats vs. low-relapsing rats, *U* = 22.5, *p* = 0.01; intermediates vs. high-relapsing rats, *U* = 55, *p* = 0.06). In contrast, DS+ discrimination ratios were similar across quartiles (Fig. 7G, intermediates vs. low-relapsing rats, *U* = 85, *p* = 0.45; high-relapsing vs. low-relapsing rats, *U* = 54, *p* = 0.35; intermediates vs. high-relapsing rats, *U* = 68, *p* = 0.17). In summary, these findings suggest that increased vigour and overall rates of cocaine self-administration in the past predicted greater responding to the DS+ during relapse testing. This highlights the critical role of the temporal pattern of drug intake in shaping subsequent relapse outcomes.

**Figure 7.**
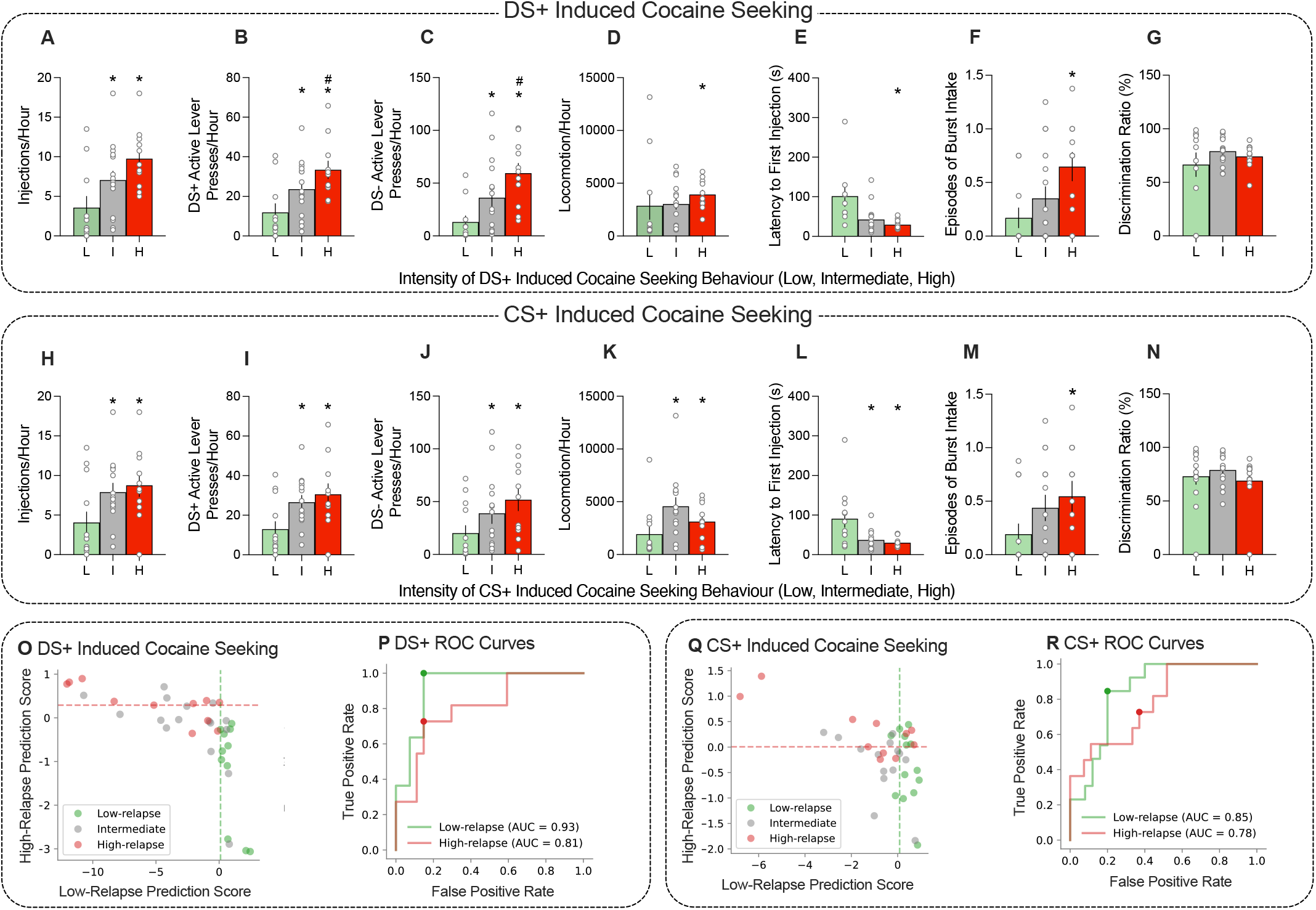
Behaviour during prior Intermittent-Access cocaine self-administration predicts the subsequent rate of cue-induced drug-seeking behaviour. Average (± SEM) **(A, H)** self-administered cocaine injections/hour, **(B, I)** active lever presses/hour during DS+ presentation, **(C, J)** active lever presses/hour during DS-presentation, **(D, K)** locomotor counts/hour, **(E, L)** latency to self-administer the first injection upon DS+ onset, **(F, M)** episodes of burst-like cocaine intake averaged across DS+ periods and **(G, N)** discrimination ratios, during the 15^th^ session of IntA training in rats showing low, intermediate or high rates of, respectively, DS+ and CS+ induced cocaine-seeking. All data are collapsed across Short-IntA and Long-IntA groups. **(O, Q)** Scatter plots representing the classification of rats based on rates of cocaine-seeking behaviour triggered by response-independent presentation of **(O)** DS+ and **(Q)** CS+. Green circles represent rats in the first quartile (low-relapse; L), red circles correspond to rats in the fourth quartile (high-relapse; H) and grey dots represent rats in the second and third quartiles (intermediate relapse; I). The dashed lines indicate the relapse thresholds for ‘high-relapse’ (red) and ‘low-relapse’ (green). Receiver operating characteristic (ROC) curves, illustrating the classification accuracy in distinguishing low-relapse and high-relapse rats based on cocaine-seeking behaviour during **(P)** DS+ and **(R)** CS+ presentations. * vs. L; #H vs. I; DS+, discriminative stimulus predicting cocaine availability; DS-, discriminative stimulus predicting cocaine unavailability; CS+, conditioned stimuli paired with cocaine delivery; For DS+ induced cocaine seeking: *n* = 11 low-relapse, *n =* 16 intermediates, *n =* 11 high-relapse; for CS+ induced cocaine seeking: *n* = 13 low-relapse, *n =* 14 intermediate relapse, *n* = 11 high-relapse.

We observed a similar pattern for CS+ responding, whereby distinct cocaine self-administration phenotypes also predicted the rate of active lever pressing during CS+ presentations. First, rats with high rates of hourly cocaine intake later responded more to the CS+ during the cocaine-seeking test (Fig. 7H; intermediates vs. low-relapsing rats, t_(25)_ = -2.21, p = 0.02; high-relapsing vs. low-relapsing rats, t_(22)_ = -2.42, p = 0.03; intermediates vs. high-relapsing rats, t_(23)_ = 0.48, p = 0.32). Second, rats with high rates of responding on the active lever during DS+ trials responded more to the CS+ during the cocaine-seeking test Fig. 7I; intermediates vs. low-relapsing rats, t_(25)_ = 2.65, p = 0.01; high-relapsing vs. low-relapsing rats, t_(22)_ = 2.74, p = 0.01; intermediates vs. high-relapsing rats, t_(23)_ = 0.65, p = 0.26). Third, rats with high rates of responding on the active lever during DS-trials also responded at higher rates to the CS+ (Fig. 7J; intermediates vs. low-relapsing rats, *U* = 50, p = 0.02; high-relapsing vs. low-relapsing rats, *U* = 27, p = 0.01; intermediates vs. high-relapsing rats, *U* = 56.5, p = 0.13). Fourth, rats with high rates of hourly locomotion showed more responding to the CS+ (Fig. 7K; intermediates vs. low-relapsing rats, *U* = 34, p = 0.01; high-relapsing vs. low-relapsing rats, *U* = 41, p = 0.04; intermediates vs. high-relapsing rats, *U* = 51, p = 0.08). Fifth, rats with the shorter latencies to initiate cocaine self-administration upon DS+ onset later responded more to the CS+ (Fig. 7L; intermediates vs. low-relapsing rats, *U* = 35, p = 0.01; high-relapsing vs. low-relapsing rats, *U* = 22, p = 0.01; intermediates vs. high-relapsing rats, *U* = 57, p = 0.32). Sixth, more episodes of burst-like cocaine intake averaged across DS+ periods predicted greater rates of responding to the CS+ (Fig. 7M, intermediates vs. low-relapsing rats, *U =* 59, *p* = 0.06; high-relapsing vs. low-relapsing rats, *U =* 38.5, *p* = 0.03; intermediates vs. high-relapsing rats, *U* = 66.5, p = 0.29). In contrast, DS+ discrimination ratios were similar across quartiles (Fig. 7N; intermediates vs. low-relapsing rats, *U* = 85, p = 0.40; high-relapsing vs. low-relapsing rats, *U* = 56.5, p = 0.20; intermediates vs. high-relapsing rats, *U* = 50, p = 0.07). Thus, increased vigour and overall rates of cocaine self-administration predict increased cocaine-seeking behaviour during subsequent CS+ presentation.

Based on the behavioural measures during IntA that defined quartile-based phenotypes, we generated a relapse prediction score for each rat to assess the extent to which differences in these measures predicted relapse propensity. Figs. 7O and Q show scatter plots used to visualize the relationship between the relapse prediction scores and our relapse stratification method during DS+ and CS+ responding, respectively. Here, each dot represents a rat colour-coded by quartile group (Green, low relapse; grey, intermediate relapse; red, high relapse). The dashed lines in Figs. 7O and Q represent classification thresholds for low relapse (green) and high relapse (red). Red dots falling above the red line were correctly predicted as showing the highest rates of cue-induced cocaine-seeking behaviour, while red dots falling below this line were falsely predicted to be “high-relapsing rats”. Similarly, green dots falling to the right of the green line were correctly predicted as showing the lowest rates of cocaine-seeking behaviour, while green dots falling to the left of this line were falsely predicted to be “low-relapsing rats”. The scatterplots in Figs. 7O and Q show that relapse prediction scores effectively distinguish between relapse phenotypes. Next, we performed a quantitative assessment of the accuracy of this classification using receiver operating characteristic curves.

Figures 7P and R illustrate receiver operating characteristic curves for DS+ and CS+ responding. The dots represent the classification threshold lines presented in the corresponding scatterplots. Dot position reflects the accuracy of the classification performance by balancing true positives and false positives. Accuracy is measured using the area under the curve (AUC) where values approaching 1.0 indicate high levels of accuracy (greater number of true positives vs. false positives) and values close to 0.5 suggest random classification. These analyses provide us with a quantitative measure to evaluate whether our method of stratification of cocaine-seeking responses during the DS+ and CS+ presentation into quartiles is reliable to predict relapse propensity. The data confirm that it is. For (DS+)-induced relapse (Fig. 7P), the receiver operating characteristic curve revealed greater AUC values for low-relapsing (0.93) compared to high-relapsing rats (0.81), indicating that our stratification method is highly reliable across these groups, especially for low-relapsing rats. For responding during the CS+ (Fig. 7R), the AUC for low-relapsing rats (0.85) was greater than that for high-relapsing rats (0.78), suggesting that our procedures can more reliably predict rats showing lower relapse scores, which is consistent with the overall low levels of responding observed during CS+ presentation. Taken together, these findings suggest that our IntA training procedure reliably predicts later propensity to seek cocaine in response to non-contingent presentations of drug-associated DSs and CSs.

## Discussion

We investigated how long (4 h/day) vs. short (2 h/day) daily bouts of intermittent cocaine self-administration influence cue-induced cocaine seeking after abstinence. Rats self-administered cocaine in the presence of drug-associated discriminative and conditioned stimuli. In both Short-IntA and Long-IntA groups, the DSs acquired control over self-administration behaviour; rats responded for cocaine during DS+ periods and suppressed responding during DS-periods. Long-IntA rats escalated their intake across sessions, ultimately consuming more than twice as much cocaine as did Short-IntA rats. After one month of forced abstinence, response-independent DS+ presentation evoked significant increases in cocaine-seeking behaviour, whereas the CS+ was ineffective. Cue-induced cocaine-seeking behaviour was similar across Short- and Long-IntA groups, indicating that with intermittent cocaine intake, cumulative intake does not predict the magnitude of cue-induced relapse. Instead, specific patterns of intake during IntA sessions—higher hourly consumption, increased lever-pressing behaviour during DS presentation, reduced latency to initiate drug self-administration as well as increased burst-like episodes —predicted relapse intensity. Thus, when cocaine self-administration is intermittent, high and escalating levels of intake are not required for increased relapse risk. Rather, individual patterns of cocaine use may more effectively mediate the neuroadaptations that drive relapse vulnerability.

### Short- and Long-IntA Cocaine Self-Administration

During IntA sessions, both Short-IntA and Long-IntA rats learned to respond for cocaine when a DS+ signalled that lever pressing would produce drug and to inhibit responding when a DS-signalled cocaine unavailability, despite continued lever access. This demonstrates that the DSs acquired strong control over drug-taking behaviour through associative learning, consistent with previous findings (Algallal et al., 2025; Ciccocioppo et al., 2001; Collins et al., 2023; Falk & Lau, 1995; Martin et al., 2022; McFarland & Ettenberg, 1997; Ndiaye et al., 2024; Pitchers et al., 2017; Weiss et al., 2000). Earlier studies using continuous-access procedures have shown that DS training during relatively brief sessions (2-3 h) can establish DS control (Ciccocioppo et al., 2001; Falk & Lau, 1995; Weiss et al., 2000). We extend these findings by showing that, under intermittent-access conditions, robust DS control over drug taking emerges even with quite limited daily DS+ exposure. Short-IntA rats experienced only 4 x 5 min of DS+ exposure/day yet still developed robust discrimination between DS+ and DS-. This highlights the potency of environmental cues in shaping drug-taking behaviour. Thus, even brief DS exposure can influence motivational states, underscoring the critical role of environmental cues in guiding drug-taking actions.

Long-IntA rats took over twice more cocaine than did Short-IntA rats. This was expected, as Long-IntA rats had twice more drug-access periods per session (8 x 5 min vs. 4 x 5 min; see also Allain & Samaha, 2019). However, when normalized for access duration, short vs. longer IntA sessions yielded remarkably similar cocaine self-administration profiles. The two groups had similar hourly rates of cocaine intake, cocaine-induced locomotor activity, lever pressing during DS+ and DS-periods, and average latency to initiate cocaine self-administration upon DS+ onset as well as average number of episodes of burst-like intake (i.e., taking > 5 injections in ≤ 5 min). Both groups also showed a pronounced drug loading effect, taking the majority of cocaine infusions during the first minute of each DS+ period, when brain cocaine concentrations would be lowest (Allain et al., 2018; Zimmer et al., 2011; Zimmer et al., 2012). This intake strategy is typically seen at the start of continuous-access drug self-administration sessions (Wilson et al., 1971; Zimmer et al., 2011) and the repeated loading each time drug becomes available is characteristic of IntA paradigms (Algallal et al., 2020; Allain et al., 2018; Collins et al., 2023; D’Ottavio et al., 2023). This volitional loading effect sensitized across sessions in both Short-IntA and Long-IntA rats. This is significant given that burst-like drug intake (i.e., taking several infusions in a short time period) is considered a behavioural marker for the development of addiction-like behaviours (Allain et al., 2015; Belin et al., 2009; Martín-García et al., 2014). In summary, under IntA conditions, session length determines total drug intake but does not influence the pattern or intensity of behavioural responding to cocaine.

Our findings also inform the long-standing association between escalation of drug intake and other addiction-relevant behaviours. Escalation remains a central focus in cocaine self-administration research (Ahmed & Koob, 1998; De Guglielmo et al., 2024; Degoulet et al., 2021), where it is often viewed as an essential feature of addiction, closely linked to, and even constitutive of, other addiction-like behaviours (De Guglielmo et al., 2024). In the present study, only Long-IntA rats escalated their cocaine intake. However, both Long- and Short-IntA rats developed a burst-like pattern of drug taking, both rapidly initiated self-administration upon signalled drug availability, and both later showed a similar intensity of cue-induced relapse to cocaine seeking. These findings suggest that, under IntA conditions, escalation of intake and the emergence of other addiction-relevant behaviours can occur independently (see also Allain et al., 2018; Rakowski et al., 2025). This challenges the widely held assumption that high and escalating levels of intake are necessary and sufficient to increase incentive motivation for the drug and promote the addiction process (see also Algallal et al., 2024; Allain et al., 2018; Allain & Samaha, 2019; D’Ottavio et al., 2023; Rakowski et al., 2025)

### Similar Cue-Induced Cocaine-Seeking Behaviour in Short- and Long-IntA Rats

Despite consuming less cocaine and experiencing fewer cue presentations, Short-IntA rats showed cue-induced cocaine seeking comparable to that seen in Long-IntA rats. This contrasts with previous studies linking greater cocaine intake to potentiated cue-induced drug seeking (Ferrario et al., 2005; Fischer et al., 2013; Jin et al., 2010; Kippin et al., 2006). Three methodological differences may account for this divergence. First, here we gave all rats a priming cocaine injection before the cue-induced drug-seeking test, such that rats experienced the cues with drug on board, as they did during prior IntA self-administration. However, we have shown previously that even without a priming injection, Long-IntA rats show the same pattern of responding to response-independent presentation of cocaine cues as observed here; increased cocaine seeking in response to the DS+, and no responding to the CS+ (Algallal et al., 2025; Ndiaye et al., 2024). Second, in prior studies, cue presentation on the relapse test day relied on the performance of drug-seeking responses and reinforced such responses (Ferrario et al., 2005; Fischer et al., 2013; Jin et al., 2010; Kippin et al., 2006). In the present study, cue presentation was response-independent. This is unlikely to explain the discrepancy, as earlier work using response-contingent CSs found that 6-8 h of continuous cocaine access achieves significantly more overall drug intake than does 6-8 h of IntA, but IntA still produces stronger CS-reinforced cocaine-seeking behaviour (James et al., 2019; Nicolas et al., 2019). Finally, prior studies used continuous-access procedures, while we used IntA. This could be a decisive difference. IntA not only more effectively models addiction-like behaviours, but also induces opposite neuroadaptations in mesolimbic dopamine signalling as compared to continuous-access procedures (Allain et al., 2015; Kawa et al., 2019; Samaha et al., 2021). Continuous access blunts cocaine-evoked dopamine responses, whereas IntA enhances them (Calipari et al., 2013, 2014). One possibility is that even limited IntA exposure (the short-IntA procedure used here) might be sufficient to cross the threshold of addiction-causing neuroplasticity, including dopamine sensitization. As this is investigated in future work, our findings align with studies showing that brief IntA exposure is as effective as extended IntA experience in promoting increased motivation to obtain drug, extinction learning and cocaine-induced reinstatement of drug seeking (Allain & Samaha, 2019). Together, the results challenge the notion of cumulative cocaine exposure as a main driver of addiction and highlight the critical role of the temporal pattern of drug use.

### Behavioural Measures Predictive of Cue-Induced Cocaine Seeking

Individual differences in behaviour during IntA self-administration predicted the intensity of cue-induced cocaine-seeking behaviour. Rats with the highest rates of responding to the DS+ or the CS+ during relapse testing were those that previously initiated cocaine self-administration most rapidly upon DS+ onset, had the highest hourly rates of drug self-administration, cocaine-induced locomotion, lever pressing during DS presentation, and number of burst-like episodes of cocaine intake. To the extent that these behaviours reflect the incentive motivational effects of cocaine (Robinson & Berridge, 1993), it would suggest that high-relapsing rats attributed greater motivational value to the drug, the drug cues, or both, making these animals more susceptible to cue-driven relapse after abstinence. Collectively, our findings support the idea that specific patterns of cocaine intake, reflecting increased incentive motivation for drug and drug cues, are stronger predictors of relapse compared to total cocaine intake (for review also see Allain et al., 2015). To the extent that our findings translate to humans, this would suggest that cocaine use patterns could be useful for identifying individuals most at risk of relapse.

## Concluding Remarks

Our findings demonstrate that the temporal pattern of cocaine intake, rather than total drug consumption, more effectively predicts vulnerability to cue-induced relapse. Even limited daily exposure to cocaine and cocaine-associated DSs under IntA conditions was sufficient to establish robust cue control over self-administration behaviour and to trigger drug-seeking actions after abstinence. High and escalating amounts of cocaine intake were not necessary for robust cue-induced drug seeking behaviour to emerge. Instead, behavioural markers indicative of heightened incentive motivation—such as rapid initiation of drug taking, increased rates of instrumental responding in the presence of DSs and a pattern of burst-like intake—were the most reliable predictors of subsequent cocaine seeking. These results challenge prevailing assumptions that total drug exposure is the main driver of addiction-related behaviours, emphasizing instead the critical influence of individual patterns of use. Understanding these patterns may improve identification of at-risk individuals and eventually inform more targeted interventions for relapse prevention.

## Acknowledgements

We thank Dr Mandy LeCocq for technical support in developing computer programmes used in this study, for advice on statistical analysis and for revising an earlier version of this manuscript. We also thank Dr. Isabel Laplante for her assistance with experimental manipulations. A.M.B is supported by a PhD merit scholarship from the Faculty of Medicine of the Université de Montréal. This work was supported by grants to A.N.S from the Canadian Institutes of Health Research (grant number 157572).

## Notes

### Competing Interest Statement

The authors have declared no competing interest.

